# Frustrated binding of biopolymer crosslinkers

**DOI:** 10.1101/483669

**Authors:** Yuval Mulla, Harmen Wierenga, Celine Alkemade, Pieter Rein ten Wolde, Gijsje H. Koenderink

## Abstract

Transiently crosslinked actin filament networks allow cells to combine elastic rigidity with the ability to deform viscoelastically. Theoretical models of semiflexible polymer networks predict that the crosslinker unbinding rate governs the timescale beyond which viscoelastic flow occurs. However a direct comparison between network and crosslinker dynamics is lacking. Here we measure the network’s stress relaxation timescale using rheology and the lifetime of bound crosslinkers using fluorescence recovery after photobleaching. lntruigingly, we observe that the crosslinker unbinding rate measured by FRAP is more than an order of magnitude slower than the rate measured by rheology. We rationalize this difference with a three-state model where crosslinkers are bound to either 0, 1 or 2 filaments, which allows us to extract crosslinker transition rates that are otherwise difficult to access. We find that the unbinding rate of singly bound crosslinkers is nearly two orders of magnitude slower than for double bound ones. We attribute the increased unbinding rate of doubly bound crosslinkers to the high stiffness of biopolymers, which frustrates crosslinker binding.

## INTRODUCTION

Cell shape and mechanics are largely governed by the actin cortex, a thin biopolymer meshwork underneath the cell membrane. The actin cortex needs to be readily deformable for processes like division and migration [1]. However the same material should resist mechanical stresses to protect the cell nucleus and plasma membrane against external stresses [2, 3]. This extraordinary combination of deformability and mechanical resistance is achieved by dynamic crosslinking proteins which stochastically bind and unbind actin filaments [4]. This design principle yields mechanical resistance as the bonds form a percolated network, whilst allowing for viscoelastic flows on timescales longer than the crosslinker unbinding timescale as the network remodels via the linker dynamics.

Rheological measurements on reconstituted actin fil-aments together with actin crosslinking proteins have shown that the microscopic crosslinker dynamics determine the macroscopic network mechanics [5-8] and mutations affecting single molecule crosslinker dynamics also change the network mechanics [9-11]. Different from most synthetic polymers, which are flexible and coil up due to thermal fluctuations, actin filaments are semiflexible polymers with a persistence length close to 10 *μ*m, on the order of the filament length [12]. Upon network deformation, the filaments are pulled taut and bending fluctuations are suppressed. The corresponding entropy reduction endows the network with an elasticity that is inversely proportional to the length scale of transverse filament fluctuations. Crosslinkers confine such fluctuations and thereby increase the stiffness of semiflexible polymer networks [13].

Whereas time-dependent mechanics of transiently crosslinked *flexible* polymer networks are well-described by a simple Maxwell model with a single stress relaxation rate [14], measurements on reconstituted actin networks [10, 15, 16] and on living cells [17-19] have revealed power law dynamics in the storage and loss modulus as a function of frequency. Theoretical modeling [20] and simulations [21] have revealed that these power law dynamics result from the superposition of multiple relaxation times of the many closely spaced crosslinkers that crosslink each filament to the surrounding network [20, 21].

Current models predict that this regime characterized by power-law dynamics occurs on timescales longer than the crosslinker unbinding time [20, 21]. Previous studies have used the onset frequency of power law dynamics to characterize the actin crosslinker *α*-actinin 4 (ACTN4) via rheological measurements and have found a crosslinker unbinding timescale on the order of 2 s [10, 16, 20]. In contrast, direct measurements of the ACTN4 binding kinetics within actin networks using Fluorescence Recovery After Photobleaching (FRAP) have found a typical unbinding time of 30 − 100 s [22-25]. Although it should be noted that these experiments have been performed under different conditions (reconstituted actin networks [10, 20] versus live cells [22-25]) the difference of more than an order of magnitude between the two sets of measurements suggests that the onset frequency of power law dynamics is not the same as the crosslinker unbinding time [20, 21].

Here we perform both FRAP and rheology experiments on reconstituted actin networks crosslinked by ACTN4. We find that the crosslinker unbinding rate measured by FRAP is an order of magnitude slower than the onset frequency of power law dynamics in the rheology even when measured on identical samples. We rationalize this difference with a three-state crosslinker model: whereas stress relaxation in the network already occurs as soon as crosslinkers unbind from one of the two filaments, full crosslinker unbinding as measured by FRAP requires detachment from both filaments. We are able to extract quantitative information on the crosslinker (un)binding kinetics which is only accessible through a combination of techniques, and has not been reported before. Interestingly, we have found that the unbinding rate of a crosslinker attached to one filament is more than an order of magnitude slower than when when it is attached to two filaments. We attribute this difference to the large bending rigidity of the actin filaments, which frustrates doubly bound crosslinkers. Our new kinetic data allow for more precise computational modeling of actin networks [21, 26], and give insight into the dynamics of the cell cortex [1, 27]. Lastly, we expect that our work will help to design synthetic materials with programmed timescales of relaxation [14, 28].

## METHODS

### Protein purification and network formation

The actin crosslinker human *α*-actinin 4 (ACTN4) was purified as described in reference [29]: Rosetta E. Coli cells were transformed to express recombinant crosslinkers with a 6xhis-tag. Induction was performed with 500 *μ*M isopropyl *β*-D-1-thiogalactopyranoside for eight hours at 25°C. After centrifugation at 6000 g for 15 minutes, cells were resuspended in 20 mM NaCl, 5 mg/ml lysozyme and 20 mM Hepes, pH 7.8. The cells were lysed by a freeze-thaw cycle, and centrifuged at 20,000 g for 30 min. The recombinant protein was purified from the supernatant using a QIAGEN nickel column. Next, the column was washed with 20-bed columns of 500 mM NaCl, 25 mM imidazole, and 20 mM Hepes, pH 7.8. The recombinant proteins were eluted with 10-bed volumes of 500 mM NaCl, 500 mM imidazole, and 20 mM Hepes, pH 7.8. The proteins were concentrated using a Centricon fil-tration device (Millipore) and purified by gel filtration in 150 mM NaCl, 20 mM Hepes pH 7.8, and 10 mM dithiothreitol (DTT) using an AKTA purifier (GE Healthcare) with a Sephadex 200 column. ACTN4 was labeled on cysteine by mixing maleimide-activated Oregon Green at a ratio of five fluorophores for every crosslinker at room temperature for 1 h. Labeled ACTN4 was separated from free dyes by gel filtration using Superdex 200 (GE Health-care) [29].

All chemicals were bought from Sigma Aldrich unless otherwise mentioned. Actin was purified from rabbit psoas skeletal muscle as described in reference [30] and stored at 80°C in G-buffer (2 mM tris-hydrochloride pH 8.0, 0.2 mM disodium adenosine triphosphate, 0.2 mM CaC*l*_2_, 0.2 mM dithiothreitol) to prevent polymerization. Unless otherwise mentioned, we used an actin concentration of 48 *μ*M, corresponding to 2 mg/ml, for all our experiments and actin was polymerized in a buffer consisting of 50 mM KCl, 20 mM imidazole pH 7.4, 2 mM MgC*l*_2_, 1 mM DTT and 0.5 mM MgATP (F-buffer). For both rheology and FRAP, all networks were allowed to polymerize at 298 K for two hours before measurements were performed. Unless otherwise mentioned, we useda crosslinker concentration of 0.48 *μ*M to obtain a molar ratio of 1/100 crosslinker/actin and on average about 1 crosslinker per 0.5 *μ*m of actin filament; under these conditions, networks are unbundled and isotropic as verified by confocal fluorescence microscopy [Fig. S4].

### Spin-down assay

A volume of 25 *μ*l monomeric (G-)actin at increasing concentrations was co-polymerized with actin binding proteins in F-buffer at room temperature, keeping the actin binding protein concentration constant (0.1 *μ*M). After two hours of polymerization the actin network together with the bound crosslinkers was spun down at 120 000 g. Afterwards, 20 *μ*l was gently pipetted from the supernatant, mixed with 20 *μ*l InstantBlue and boiled at 95°C for 5 minutes in a closed Eppendorf vial. 30 *μ*l of this solution was loaded onto a 4-15 % Mini-PROTEAN TGX™ Precast Protein Gel with 10 wells of 30 μl, purchased at Bio-Rad. Gels were run for 30 minutes at 200 V, washed with Mili-q water, stained overnight with InstantBlue and washed three times with tap water. Band intensities of the ACTN4 in the supernatant were quantified using ImageJ. The fraction of bound linkers *φ_bound_* was determined by subtracting and normalizing the ACTN4 band intensity at a particular actin concentration by the band intensity in the absence of actin: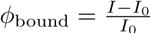. Background correction was applied to all band intensities by subtracting the average intensity of a region adjacent to the band of interest.

### Fluorescence Recovery After Photobleaching

The bound crosslinker lifetime within actin networks was measured via Fluorescence Recovery After Photobleaching (FRAP) using a Nikon A1 confocal microscope with a perfect focus system, a 100x oil immersion objective, and a 100-mW 488 nm argon ion laser. The temperature was controlled by a home-built temperature regulator. The regulator consisted of a temperature-controlled water bath connected to both a home-made objective heater and a sample glass slide heater. The temperature of the water was measured inside the objective heater via a P1000 temperature sensor. We calibrated the temperature inside the sample, measured by inserting a 0.025 mm thermocouple in a glow channel filled with deionized water, against the temperature inside the objective heater. Using this set-up, temperatures between 285−333 K can be achieved.

The FRAP protocol started with 10 images to determine baseline fluorescence. Next, bleaching was performed by using a high intensity laser power such that 50 − 70 % of the fluorescence intensity was bleached in0.5 seconds. The fluorescence recovery was tracked during a period of approximately 5x the typical recovery time, with a sampling rate that halved every 10 frames,starting with 10 frames per second. Unless otherwise mentioned, a circular area was bleached of 2 *μ*m radius and an equally sized area was used as a reference. The laser intensity during imaging was chosen such that the reference intensity dropped less than 5 % during the recovery phase. To extract a recovery rate, the normalized intensity during recovery *I/I*_ref_ was fitted with a single exponent *I/I*_ref_ =1-*I*_0_*/I*_ref_ exp(-*t k_FRAP_*), where *I*_0_ is the intensity directly after bleaching and *k*_FRAP_ the recovery rate. The timescale of recovery is governed by the typical crosslinker diffusion time, which scales quadratically with the bleach radius, and the typical crosslinker unbinding time, which is independent of the bleach radius. To dissect these two contributions, we compared the recovery time for different bleach radii, 2 *μ*m and 4 *μ*m. We did not find a statistically significant difference [Fig. S4]. This result is consistent with a calculation where we estimate that the typical diffusion distance in the timescale of FRAP recovery time is significantly larger than the FRAP radius (40 *μ*m vs. 2 *μ*m). We used a diffusion constant based on the Stokes-Einstein relationship, assuming a crosslinker radius of 3 nm on basis of the crystal structure [31].

### Rheology

Rheology was performed using a stress-controlled Kinexus Malvern Pro rheometer with a stainless steel 20 mm radius cone plate geometry with a °egree an-gle. We loaded 40 *μ*l G-actin mixed with ACTN4 directly after mixing the proteins into the polymerization buffer (F-buffer). A thin layer of Fluka mineral oil Type A was added around the geometry to prevent evaporation, and the sample was closed off with a hood to minimize effects of air flow. Polymerization of the network was followed by applying a small oscillatory shear with a strain amplitude of 0.5 % and a frequency of 0.5 Hz for 2 h. After 2 h of polymerization, a frequency sweep was performed between 0.01 − 10 Hz, using 10 data points per decade. Frequencies above 10 Hz could not be measured as inertial effects from the rheometer dominated the rheological response of the actin network at high frequencies. After characterization at 25°C (298 K), the temperature was adjusted and equilibrated for 10 minutes. The typical frequency was extracted from the frequency dependent storage and loss moduli (*G′* and *G″*) by fitting a previously published model which considers the multi-relaxation times due to many crosslinks unbinding per filament [20]. The model is based on the nonlinear-force extension curve of a semiflexble filament, and uses mean-field arguments to extract mechanical properties of the network from the single filament fluctuations:

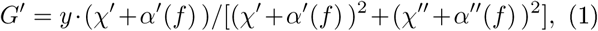

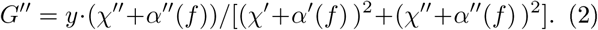

*y* is a prefactor to allow for direct comparison between the semi-quantitative model and experimental data, while *χ* describes the viscous drag limiting transverse filament fluctuations:

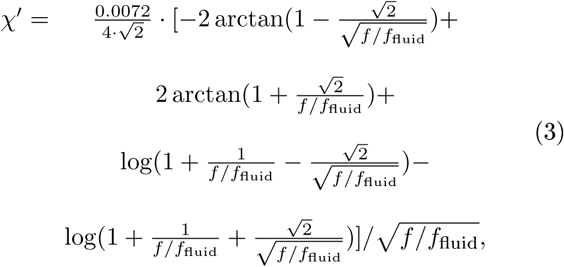

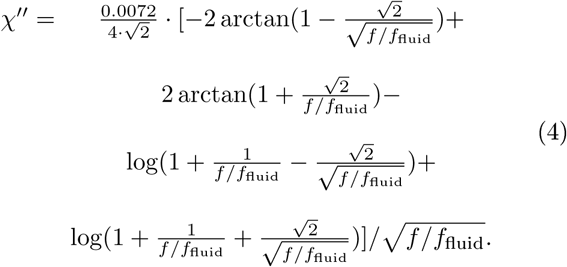

Here *f*_fluid_ is the timescale of the fluid drag which depends on the fluid viscosity and the mesh size [32] and is typically on the order of 100 Hz for actin networks [33]. Lastly, *α* describes the effect of the crosslinkers limiting transverse filament fluctuations:

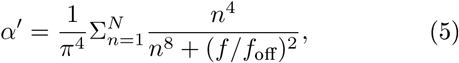

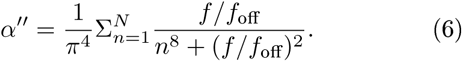

Here *N* is the number of crosslinkers per filament and *f*_off_ is the off-rate of the crosslinker. In this work, we refer to this rate as the rheology rate, *k*_rheo_, to prevent confusion with the FRAP rate *k*_FRAP_, which is also governed by crosslinker unbinding. The *G′*~*G″*~*f*^−1/2^power law extends down to lower frequencies (i.e. larger time scales) for higher numbers of crosslinkers per filament. In line with previous research [20], we arbitrarily assume the number of crosslinkers (*N*) to be 10 per filament, which is likely smaller than the actual number of crosslinkers per filament but still large enough to observe power law behavior of the shear moduli over the full experimental regime. Figure S2 contains the frequency sweeps over the full range of measured temperatures.

## RESULTS

We measure the crosslinker unbinding rate in actin networks using Fluorescence Recovery After Photobleaching (FRAP) by bleaching a circle of 2 *µ*m radius using a high intensity laser power. Afterwards, we track the recovery of fluorescent intensity within this circle and use a reference area to correct for any photobleaching during imaging of the recovery phase. We find that the fluorescent intensity recovery is well-described with a single exponential function with a rate of *k*_FRAP_ = 0.036 ±0.001 s^−1^, and that the intensity asymptotically approaches the fluorescent intensity before bleaching, indicating that all crosslinkers are mobile [Fig. 1a].

**Figure 1.**
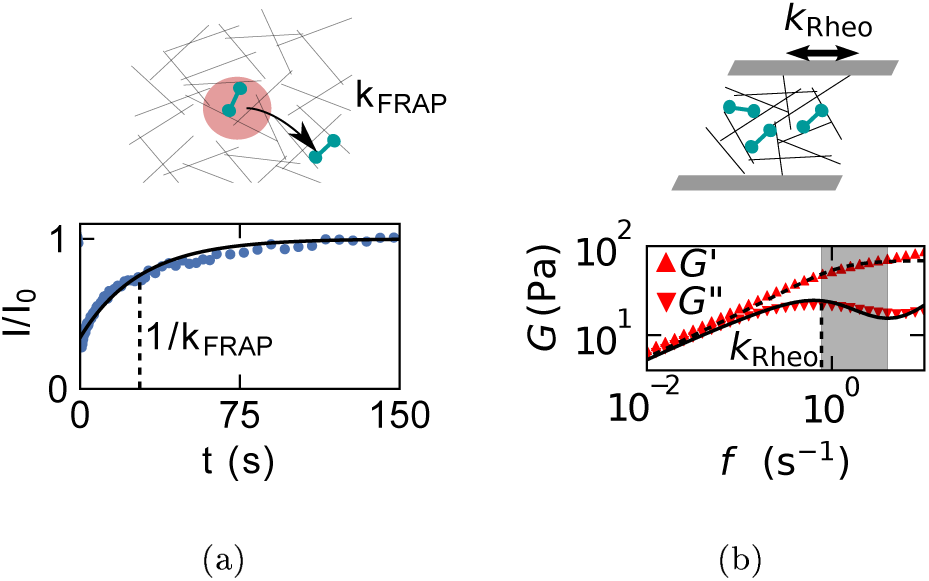
Measurements of the FRAP and rheology rates. a) FRAP curve of the fluorescence intensity as function of time after photobleaching (blue dots) together with a fit to a mono-exponential recovery (black line) with a rate of *k*_FRAP_ = 0.036 0.001 s^−1^ (dashed line, standard error on basis of 4 repeats). b) Frequency dependence of the storage and loss moduli (see legend) together with a fit to the crosslink governed networks dynamics model (dashed and solid lines, Eq. 1 and 2). *G″* peaks at *k*_Rheo_ = 0.77 0.03 s^−1^ (vertical dashed line, standard error on basis of fit). The shading de-marcates the intermediate frequency regime from the low and high frequency regimes. Both FRAP and rheology data are obtained at 283 K. The actin filaments (black), crosslinkers (green) and FRAP area (red) in the schematics are not drawn to scale.

Next, we measure network mechanics using small amplitude oscillatory shear rheology: we apply an oscillatory strain of 0.5 % amplitude and measure the stress required for this deformation. We analyze the in-phase and out-of-phase contributions (storage and loss component respectively) as a function of the applied frequency. Consistent with earlier observations on transiently cross linked actin networks [8, 10, 15], we observe that the mechanical response can be divided into three frequency regimes [Fig. 1b]. Firstly, at low frequencies we observe power law dynamics, consistent with a recent crosslink-governed network dynamics model which predicts a *G′*~*G″*~*f*^−1/2^scaling at frequencies below the crosslinker unbinding rate [20]. Secondly, at intermediate frequencies, the storage modulus increases less steeply while the loss modulus decreases as crosslinker unbinding becomes increasingly unlikely [13, 20]. Lastly, at high frequencies the storage and loss modulus increase again as transverse filament fluctuations are hampered by viscous drag of the surrounding fluid [33]. We note that there is in principle also a fourth regime at very low frequencies, where filaments exhibit terminal relaxation on timescales long enough to allow for filament relaxation over its full length. However, this regime is beyond the accessible timescales.

We fit our experimental data over all three regimes using the cross-link governed network dynamics model [20],giving a rate *k*_Rheo_ of 0.77± 0.03 s^−1^ (Eq. 2, indicated by the vertical line in Fig. 1b). The only other fit parameters are the timescale of fluid drag (*ω* _fluid_ = 45 s^−1^) and a pre-factor to scale both *G′*and *G*” (*y* = 0.02 Pa) as the model is only semi-quantitative [20]. According to the model [20], *k*_Rheo_ should correspond to the inverse of to the crosslinker unbinding timescale, yet the FRAP rate is more than an order of magnitude slower at the same temperature.

To test the robustness of this difference in rates, we perform FRAP and rheology measurements as a function of temperature (280-298 K). As shown in Fig. 2, the FRAP and rheology rates both increase with temperature in accordance with the Arrhenius equation for thermally activated processes, *k*(*T*)~exp(*-E*_A_*/T*), where *k* is the rate, *T* is the temperature. We can therefore extract activation energies *E*A for the FRAP rate of 33 ±4 *k*_B_*T* and for the rheology rate of 75 ±3 *k*_B_*T* [Fig. 2]. Interestingly, this implies that the onset frequency for stress relaxation as measured by rheology is significantly more temperature-dependent than the crosslinker unbinding rate as measured by FRAP. Furthermore, the onset frequency for stress relaxation, *k*_*rheo*_, is higher than the crosslinker unbinding rate over the full range of measured temperatures.

**Figure 2.**
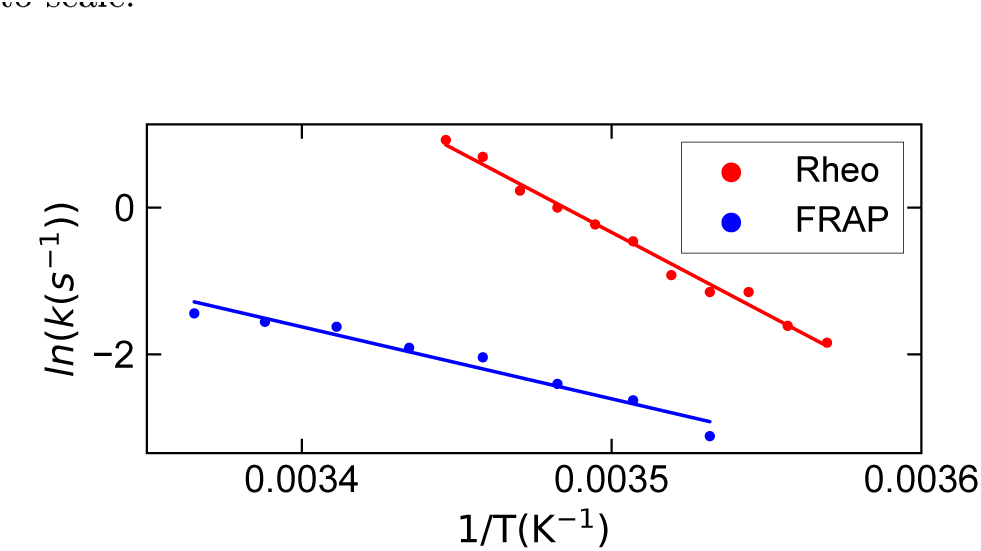
Arrhenius plot of the characteristic crosslink unbinding rates inferred from FRAP and rheology analysis. Fitting both sets of data to the Arrhenius equation (solid lines) yields an activation energy of 33±4 *k*_B_ *T* for FRAPand 75 ± 3 *k*_B_ *T* for rheology.

In order to explain the ~20-fold difference in timescale between the characteristic rates inferred from FRAP and rheology, we hypothesize that stress relaxation in crosslinked actin networks occurs as soon as one of the actin-binding domains of a crosslinker unbinds. In contrast, crosslinker redistribution as measured by FRAP requires both binding domains to detach. To formalize this hypothesis, we model the crosslinker dynamics by a three-state model where crosslinkers are either bound to zero (*s*_0_), one (*s*_1_) or two filaments (*s*_2_). In this framework, the onset frequency for stress relaxation is defined by the rate of an *s*2 crosslinker to unbind from one filament:

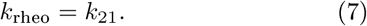

The FRAP rate is more complex, as it requires unbinding from both filaments. Note that this rate is potentially much longer than the rate *k*_21_as measured by rheology, as cycling between the singly and doubly bound state can occur many times before the crosslinker fully unbinds. We hypothesize that this cycling between both binding sites explains the difference in timescales between FRAP and rheology observed experimentally. As explained in detail in the Supplementary Information, we find an analytical solution for the asymptotic rate at which singly bound crosslinkers (*s*_1_) and doubly bound crosslinkers (*s*_2_) make a transition to the fully unbound state (*s*_0_):

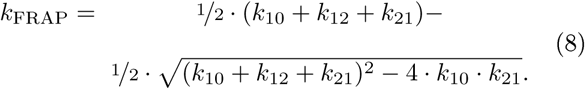

The FRAP rate contains two unknown parameters, *k*_10_ and *k*_12_, which we can dissect by measuring the recovery rate as a function of the actin concentration: whereas the unbinding rates *k*_10_ and *k*_21_ are independent of the actin concentration, the binding rate *k*_12_ increases linearly as the average filament-filament distance decreases. Assuming that there is no spatial correlation between filaments:

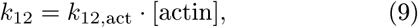

where [actin] is the concentration of actin in *μ*M and *k*_12,act_ the binding rate at an actin concentration of 1 *μ*M. Consequently, the three-state model predicts that the FRAP rate decreases with an increase of the actin concentration, as crosslinkers reside more often in the doubly bound state. Consistent with this prediction, we find that *k*_FRAP_ decreases from 0.31±0.04 ^−1^at 24 *μ*M actin to 0.21 ±0.02 s^−1^at 60 *μ*M actin [Fig. 3a]. In contrast, the onset frequency of stress relaxation as measured with rheology does not depend on the actin concentration (Refs. [8, 15, 20] and Fig. S5), as the unbinding rate *k*_21_ is independent from the actin concentration. We next combine eq. 8 and 9 to fit the FRAP data as a function of the actin concentration and find 0.43±0.06 s^−1^for the unbinding rate *k*_10_ and 0.3±0.1 s^−1^ for the binding rate *k*_12_ at an actin concentration of 48 *μ*M.

**Figure 3.**
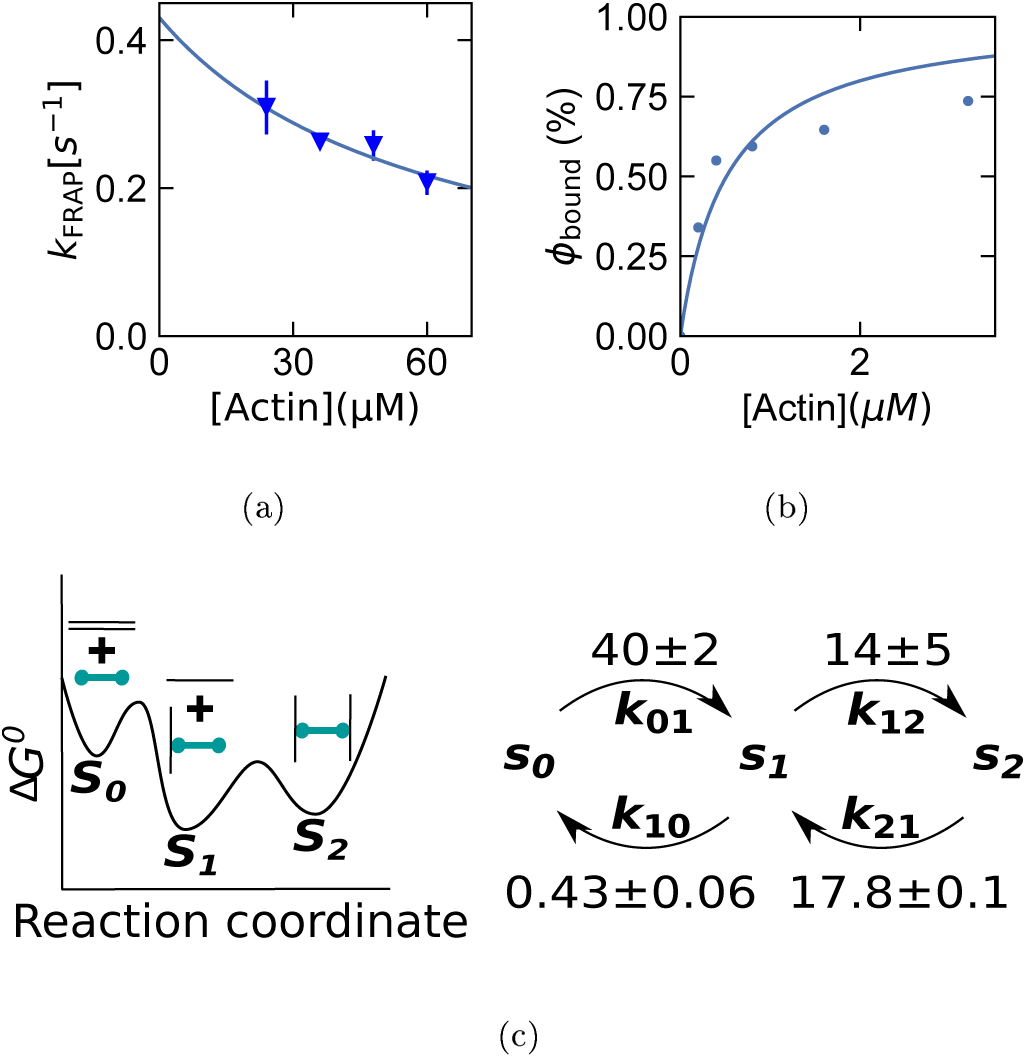
Retrieving crosslinker binding and unbinding rates from a three-state model. a) The FRAP rate as a function of actin concentration measured at 298 K is fitted with the three-state model (line; eqs. 8 and 9) to extract *k*_10_ and *k*_12_. The error bars are the standard error calculated on basis of 4 repeats per condition. b) The fraction of bound crosslinkers as a function of the actin concentration is fitted with Eq. 11 (line) to extract *k*_01_. c) All rates are in *s*^−1^ and with standard error on basis of the fit, measured at 25°C at an actin concentration of 48 *μ*M. The rate *k*_21_ is extrapolated to *T* = 25°C using the Arrhenius plot of the onset frequency of stress relaxation [Fig. 2].

Lastly, in order to measure the rate at which an unbound crosslinker binds to a filament, *k*_01_, we perform a spindown assay to separate actin-bound crosslinkers (*s*_1_ + *s*_2_) from freely diffusing crosslinkers (*s*_0_) as a function of the actin concentration. We find that the fraction of bound crosslinkers increases with the concentration of actin [Fig. 3b]. Like *k*_12_, the binding rate *k*_01_ increases linearly with the actin concentration:

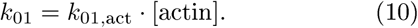

Due to detailed balance, we know that the fraction of bound linkers depends on all (un)binding rates according to (see SI for the derivation):

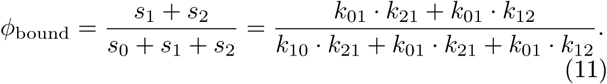

Using this equation, we fit the spin-down data to extract the last unknown rate, *k*_01,act_ and find 0.83 0.03 s^−1^. All rates are summarized in Fig. 3c.

## DISCUSSION

We have compared the dynamics of crosslinker unbinding in actin networks as measured by fluorescence recovery after photobleaching and by rheology. Surprisingly, we have found that the crosslinker unbinding rate measured by FRAP is more than an order of magnitude slower than the onset frequency of stress relaxation measured by rheometry. We have rationalized this difference with a three-state model where crosslinkers are bound to either 0, 1 or 2 filaments. Whereas stress relaxation in the network already occurs as soon as crosslinkers unbind from one of the two filaments, full crosslinker unbinding as measured by FRAP requires detachment from both filaments. Our results are consistent with rheology [10, 20] and FRAP measurements [22-25] in literature and provide a mechanistic explanation for the difference in rates. Furthermore, we have used this model to extract crosslinker transition rates that are otherwise difficult to access.

Interestingly, we have found that the unbinding rate of a crosslinker attached to one filament is more than an order of magnitude slower than when two filaments are attached (*k*_10_ = 0.43± 0.06 s^−1^ vs. *k*_21_ = 17.8 ±0.1 s^−1^). This difference suggests that binding of a second filament causes crosslinker frustration, which speeds up crosslinker-filament dissociation. This crosslinker frustration is different from the frustration of actin filaments in crosslinked bundles [34], which is due to the helical pitch of actin filaments [35, 36]. Instead, we attribute the increased crosslinker unbinding to deformation of the crosslinker [37, 38]. Steered molecular dynamics have revealed that torsion in the ACTN4 backbone is energetically highly unfavorable and a 180° torsion requires ~75 *k*_B_*T*, and similarly a crosslinker extension of 10 % requires ~70 *k*_B_*T* [37]. As the persistence length of actin filaments is large (~10 *μ*m [12]) compared to the typical crosslinker distance (~0.5 *μ*m), binding of two filaments likely causes constraints in the crosslinker orientation and length. Therefore, we speculate that crosslinker frustration should be common in crosslinked semiflexible polymer networks. Our work provides an experimentally straightforward way to characterize molecular binding kinetics of crosslinkers within networks by combining rheology with FRAP measurements, for example as a function of the crosslinker compliance [39, 40].

## Supporting information

## ACKNOWLEDGMENTS

We thank Wouter Ellenbroek, Nicholas Tito, Fred MacKintosh and Chase Broedersz for fruitful discussions. We thank William Brieher and Vivian Tang for the kind gift of purified ACTN4, Marjolein Kuit-Vinkenoog for actin purification, Henk-Jan Boluit for design and Philippine Doekes for testing and calibrating the temperature control. This work is part of the research program of the Netherlands Organisation for Scientific Research (NWO). We gratefully acknowledge financial support from an ERC Starting Grant (335672-MINICELL) and from an ERC Synergy grant (609822-MODEL CELL).

## REFERENCES

[1] G. Salbreux, G. Charras, and E. Paluch. Actin cortex mechanics and cellular morphogenesis. Trends in cell biology, 22(10):536–45, oct 2012.

[2] J. Y. Tinevez, U. Schulze, G. Salbreux, J. Roensch, J.-F. Joanny, and Ewa Paluch. Role of cortical tension in bleb growth. Proceedings of the National Academy of Sciences, 106(44):18581–18586, nov 2009.

[3] C. Denais, R. M. Gilbert, p. lsermann, A. L. McGregor, M. te Lindert, B. Weigelin, p. M. Davidson, p. Friedl, K. Wolf, and J. Lammerding. Nuclear envelope rupture and repair during cancer cell migration. Science, 352(6283):353–358, apr 2016.

[4] T. D. Pollard. Actin and Actin-Binding Proteins. Cold Spring Harbor Perspectives in Biology, 8(8):a018226, aug 2016.

[5] D.H. Wachsstock, W.H. Schwarz, and T.D. Pollard. Cross-linker dynamics determine the mechanical properties of actin gels. Biophysical Journal, 66(3):801–809, mar 1994.

[6] J. Xu, D. Wirtz, and T. D. Pollard. Dynamic Cross-linking by *a*-Actinin Determines the Mechanical Properties of Actin Filament Networks. Journal of Biological Chemistry, 273(16):9570–9576, apr 1998.

[7] R. Tharmann, M. M. A. E. Claessens, and A. R. Bausch. Viscoelasticity of isotropically cross-linked actin networks. Physical Review Letters, 98(8):8–11, 2007.

[8] O. Lieleg, K. M. Schmoller, M. M. A. E. Claessens, and A. R. Bausch. Cytoskeletal polymer networks: Viscoelastic properties are determined by the microscopic interaction potential of cross-links. Biophysical Journal, 96(11):4725–4732, 2009.

[9] A. Weins, J. S. Schlondorff, F. Nakamura, B. M. Denker, J. H. Hartwig, T. P. Stossel, and M. R. Pollak. Disease-associated mutant alpha-actinin-4 reveals a mechanism for regulating its F-actin-binding affinity. Proceedings of the National Academy of Sciences of the United States of America, 104(41):16080–5, 2007.

[10] N. Y. Yao, D. J. Becker, C. P. Broedersz, M. Depken, F. C. MacKintosh, M. R. Pollak, and D. A. Weitz. Non-linear Viscoelasticity of Actin Transiently Cross-linked with Mutant *α*-Actinin-4. Journal of Molecular Biology, 411(5):1062–1071, sep 2011.

[11] M. Maier, K. W. Müller, C. Heussinger, S. Köhler, W. A. Wall, A. R. Bausch, and O. Lieleg. A single charge in the actin binding domain of fascin can independently tune the linear and non-linear response of an actin bundle network. The European Physical Journal E, 38(5):50, 2015.

[12] F. Gittes, B Mickey, J Nettleton, and J Howard. Flexural rigidity of microtubules and actin filaments measured from thermal fluctuations in shape. The Journal of cell biology, 120(4):923–34, feb 1993.

[13] F. C. MacKintosh, J. Käs, and P. A. Janmey. Elasticity of Semiflexible Biopolymer Networks. Physical Review Letters, 75(24):4425–4428, dec 1995.

[14] S. C. Grindy, M. Lenz, and N. Holten-Andersen. Engineering Elasticity and Relaxation Time in Metal-Coordinate Cross-Linked Hydrogels. Macromolecules, 49(21):8306–8312, nov 2016.

[15] O. Lieleg, M. M. A. E. Claessens, Y. Luan, and A. R. Bausch. Transient binding and dissipation in cross-linked actin networks. Physical Review Letters, 101(10):1–4, 2008.

[16] Y. Mulla, F. C. MacKintosh, and G. H. Koenderink. Origin of soft glassy rheology in the cytoskeleton. Arxiv, pages 1-6, oct 2018.

[17] B. Fabry, G. N. Maksym, J. P. Butler, M. Glogauer, D. Navajas, and J. J. Fredberg. Scaling the Microrheology of Living Cells. Physical Review Letters, 87(14):148102, sep 2001.

[18] X. Trepat, G. Lenormand, and J. J. Fredberg. Universality in cell mechanics. Soft Matter, 4(9):1750, 2008.

[19] E. Fischer-Friedrich, Y. Toyoda, C. J. Cattin, D. J. Müller, A. A. Hyman, and F. Jülicher. Rheology of the Active Cell Cortex in Mitosis. Biophysical Journal, 111(3):589–600, aug 2016.

[20] C. P. Broedersz, M. Depken, N. Y. Yao, M. R. Pollak, D. A. Weitz, and F. C. MacKintosh. Cross-Link-Governed Dynamics of Biopolymer Networks. Physical Review Letters, 105(23):238101, nov 2010.

[21] K. W. Müller, R. F. Bruinsma, O. Lieleg, A. R. Bausch, W. A. Wall, and A. J. Levine. Rheology of semiflexible bundle networks with transient linkers. Physical Review Letters, 112(23):1–5, 2014.

[22] P. Hotulainen and P. Lappalainen. Stress fibers are generated by two distinct actin assembly mechanisms in motile cells. The Journal of Cell Biology, 173(3):383–394, may 2006.

[23] J. R. Michaud, M. Hosseini-Abardeh, K. Farah, and C. R. J. Kennedy. Modulating *α*-actinin-4 dynamics in podocytes. Cell Motility and the Cytoskeleton, 66(3):166– 178, mar 2009.

[24] A. J. Ehrlicher, R. Krishnan, M. Guo, C. M. Bidan, D. A. Weitz, and M. R. Pollak. Alpha-actinin binding kinetics modulate cellular dynamics and force generation. Proceedings of the National Academy of Sciences, 112(21):201505652, 2015.

[25] E. S. Schiffhauer, T. Luo, K. Mohan, V. Srivastava, X. Qian, E. R. Griffis, p. A. Iglesias, and D. N. Robinson. Mechanoaccumulative Elements of the Mammalian Actin Cytoskeleton. Current Biology, 26(11):1473–1479, jun 2016.

[26] V. Wollrab, J. M. Belmonte, M. Leptin, F. Nedelec, and G. H. Koenderink. Polarity sorting drives remodeling of actin-myosin networks. bioRxiv, 2018.

[27] Y. Mulla and G. H. Koenderink. Crosslinker mobility weakens transient polymer networks. Arxiv, may 2018.

[28] N. B. Tito, C. Creton, C. Storm, and W. G. Ellenbroek. Harnessing entropy to enhance toughness in reversibly crosslinked polymer networks. Arxiv, pages 1-19, oct 2018.

[29] V. W. Tang and W. M. Brieher. alpha-Actinin-4/FSGS1 is required for Arp2/3-dependent actin assembly at the adherens junction. Journal of Cell Biology, 196(1):115– 130, 2012.

[30] J. Alvarado, M. Sheinman, A. Sharma, F. C. MacKintosh, and G. H. Koenderink. Molecular motors robustly drive active gels to a critically connected state. Nature Physics, 9(9):591–597, sep 2013.

[31] E. s De Almeida Ribeiro, N. Pinotsis, A. Ghisleni, A. Salmazo, P. V. Konarev, J. Kostan, B. Sjöblom, C. Schreiner, A. A. Polyansky, E. A. Gkougkoulia, M. R. Holt, F. L. Aachmann, B. Zagrovic, E. Bordignon, K. F. Pirker, D. I. Svergun, M. Gautel, and K. Djinovic-Carugo. The structure and regulation of human muscle *α*-Actinin. Cell, 159(6):1447–1460, 2014.

[32] H. I C. G. de Cagny, B. E. Vos, M. Vahabi, N. A. Kurniawan, M. Doi, G. H. Koenderink, F. C. MacKintosh, and D. Bonn. Porosity Governs Normal Stresses in Polymer Gels. Physical Review Letters, 117(21):217802, nov 2016.

[33] G. H. Koenderink, M. Atakhorrami, F. C. MacKintosh, and C. F. Schmidt. High-frequency stress relaxation in semiflexible polymer solutions and networks. Physical Review Letters, 96(13):1–4, 2006.

[34] H. Shin, K. R. Purdy Drew, J. R. Bartles, G. C. L. Wong, and G. M. Grason. Cooperativity and Frustration in Protein-Mediated Parallel Actin Bundles. Physical Review Letters, 103(23):238102, nov 2009.

[35] C. Heussinger, M. Bathe, and E. Frey. Statistical Mechanics of Semiflexible Bundles of Wormlike Polymer Chains. Physical Review Letters, 99(4):048101, jul 2007.

[36] M. M. A. E. Claessens, C. Semmrich, L. Ramos, and A. R. Bausch. Helical twist controls the thickness of F-actin bundles. Proceedings of the National Academy of Sciences, 105(26):8819–8822, jul 2008.

[37] J. Golji, R. Collins, and M. R. K. Mofrad. Molecular Mechanics of the *α*-Actinin Rod Domain: Bending, Torsional, and Extensional Behavior. PLoS Computational Biology, 5(5):e1000389, may 2009.

[38] D. S. Courson and R. S. Rock. Actin cross-link assembly and disassembly mechanics for alpha-Actinin and fascin. The Journal of biological chemistry, 285(34):26350–7, aug 2010.

[39] M. L. Gardel, F. Nakamura, J. H. Hartwig, J. C. Crocker, T. P. Stossel, and D. A. Weitz. Prestressed F-actin networks cross-linked by hinged filamins replicate mechanical properties of cells. Proceedings of the National Academy of Sciences of the United States of America, 103(6):1762–1767, 2006.

[40] K.E. Kasza, F. Nakamura, S. Hu, p. Kollmannsberger, N. Bonakdar, B. Fabry, T.P. Stossel, N. Wang, and D.A. Weitz. Filamin A Is Essential for Active Cell Stiffening but not Passive Stiffening under External Force. Biophysical Journal, 96(10):4326–4335, may 2009.

