## Supplementary material for "Frustrated binding of biopolymer crosslinkers"

### Frustrated binding of biopolymer crosslinkers - Supplementary Information

Yuval Mulla[1], Harmen Wierenga[1], Celine Alkemade[1,2], Pieter Rein ten Wolde[1], Gijsje H. Koenderink[1*]
[1]*Living Matter Department, AMOLF, Science Park 104, 1098 XG Amsterdam and*
[2]*Department of Bionanoscience, Faculty of Applied Sciences,*
*Delft University of Technology, Delft, Netherlands*

#### FIRST PASSAGE TIME DISTRIBUTION IN THREE STATE MODEL

We are interested in the rate at which actin-binding crosslinkers fully unbind. We consider crosslinkers with two 'heads', i.e. binding sites for actin filaments. This means they can be in three states; unbound, single head bound, or two heads bound. Bound crosslinkers start off in a distribution over both states 1 (single head bound) and 2 (both heads bound). Here, we will calculate the first passage time distributions to the full unbinding (state 0), given that the crosslinkers start in state 1 or state 2. The combined first passage time distribution is a linear combination of these two subdistributions. We will show that this is a double exponential distribution, but that it has a single exponential tail. Furthermore, we will demonstrate that the FRAP recovery curve is governed by this tail, and we calculate the effective unbinding rate in terms of the crosslinker (un)binding rates.

We denote the first passage time distribution from state $i$ to state $j$ by $f_{i,j}(t)$, which gives the probability density that a crosslinker starting at state $i$ reaches state $j \neq i$ for the first time after a period $t$. Hence, our goal is to calculate $f_{1,0}(t)$ and $f_{2,0}(t)$. The latter distribution can be decomposed into the first time $t'$ that the crosslinker arrives from state 2 to state 1, and the time $t-t'$ it takes to finally make it to state 0,

$$f_{2,0}(t) = \int_0^t f_{2,1}(t')\, f_{1,0}(t-t')\,\mathrm{d}t'. \tag{1}$$

This convolution can be resolved using the Laplace transform

$$\widetilde{f}_{i,j}(s) \equiv \int_0^\infty f_{i,j}(t)\,\mathrm{e}^{-st}\,\mathrm{d}t, \tag{2}$$

which changes the convolution of Eq. 1 into a simple product

$$\widetilde{f}_{2,0}(s) = \widetilde{f}_{2,1}(s)\,\widetilde{f}_{1,0}(s)\,. \tag{3}$$

There is only a single rate leaving from state 2, giving a simple form of the first passage time distribution to state 1:

$$f_{2,1}(t) = k_{21}\mathrm{e}^{-k_{21}t}, \tag{4}$$

where $k_{21}$ is the rate at which crosslinkers go from state 2 to state 1. The Laplace transform of this equation gives

$$\widetilde{f}_{2,1}(s) = \frac{k_{21}}{s+k_{21}}. \tag{5}$$

There are two options for a crosslinker bound with a single head, namely (re)binding of the unbound head or complete unbinding. This means that the first passage time probability $f_{1,0}(t)$ can be split into a part where the first step is unbinding, and a part where the first step involves full binding. When the first step is towards state 2 at time $t'$, then a first passage is required from state 2 to state 0 in the remaining time, leading to

$$\begin{aligned} f_{1,0}(t) =& k_{10}\mathrm{e}^{-(k_{10}+k_{12})t} \\ &+ \int_0^t k_{12}\mathrm{e}^{-(k_{10}+k_{12})t'} f_{2,0}(t-t')\,\mathrm{d}t'. \end{aligned} \tag{6}$$

Again, we perform a Laplace transform on both sides of the equation to resolve the convolution. Eq. 3 and Eq. 5 subsequently are used to create a linear equation for $\widetilde{f}_{1,0}(s)$,

$$\begin{aligned} \widetilde{f}_{1,0}(s) =& \frac{k_{10}}{s+k_{10}+k_{12}} \\ &+ \frac{k_{12}}{s+k_{10}+k_{12}}\frac{k_{21}}{s+k_{21}}\widetilde{f}_{1,0}(s)\,. \end{aligned} \tag{7}$$

The solution of this equation is given by

$$\widetilde{f}_{1,0}(s) = \frac{k_{10}(s+k_{21})}{s^2+(k_{10}+k_{12}+k_{21})\,s+k_{10}k_{21}} \tag{8}$$

$$= \frac{k_{10}}{r_2-r_1}\left[\frac{k_{21}-r_1}{s+r_1}+\frac{r_2-k_{21}}{s+r_2}\right], \tag{9}$$

while through Eq. 3,

$$\widetilde{f}_{2,0}(s) = \frac{k_{10}k_{21}}{s^2+(k_{10}+k_{12}+k_{21})\,s+k_{10}k_{21}} \tag{10}$$

$$= \frac{k_{10}k_{21}}{r_2-r_1}\left[\frac{1}{s+r_1}-\frac{1}{s+r_2}\right]. \tag{11}$$

Here, we defined the two positive effective rates

$$\begin{aligned} r_1 =& \frac{1}{2}(k_{10}+k_{12}+k_{21}) \\ &- \frac{1}{2}\sqrt{(k_{10}+k_{12}+k_{21})^2-4k_{10}k_{21}}, \end{aligned} \tag{12}$$

$$\begin{aligned} r_2 =& \frac{1}{2}(k_{10}+k_{12}+k_{21}) \\ &+ \frac{1}{2}\sqrt{(k_{10}+k_{12}+k_{21})^2-4k_{10}k_{21}}. \end{aligned} \tag{13}$$

For a general initial state, of which the fraction in state 1 is $\alpha$, and thus the fraction in state 2 is $(1-\alpha)$, the first passage time distribution from that general state to state 0 is defined as

$$f_{2-\alpha,0}(t) = \alpha f_{1,0}(t) + (1-\alpha)\, f_{2,0}(t) \tag{14}$$

The general solution can be found by applying inverse Laplace transforms to Eq. 9 and Eq. 11. We solve the complex integrals using Cauchy's residue theorem, which gives

$$\begin{aligned} f_{2-\alpha,0}(t) =& \frac{k_{10}}{r_2 - r_1} \\ & \times \left[ (k_{21} - \alpha r_1)\, \mathrm{e}^{-r_1 t} - (k_{21} - \alpha r_2)\, \mathrm{e}^{-r_2} \right] . \end{aligned} \tag{15}$$

This is the general solution for the first passage time distribution. It can be shown that $r_1 < k_{21} < r_2$, and since $0 \le \alpha \le 1$, we may write

$$\begin{aligned} f_{2-\alpha,0}(t) =& \frac{k_{10}}{r_2 - r_1} (k_{21} - \alpha r_1)\, \mathrm{e}^{-r_1 t} \\ & \times \left[ 1 - \frac{k_{21} - \alpha r_2}{k_{21} - \alpha r_1} \mathrm{e}^{-(r_2 - r_1)} \right] . \end{aligned} \tag{16}$$

This distribution contains an exponential tail with a rate $k_{tail}$ which equals

$$k_{tail} = \lim_{t\to\infty} \frac{\log\left[ f_{2-\alpha,0}(t) \right]}{t} = \lim_{t\to\infty} \frac{\dot{f}_{2-\alpha,0}(t)}{f_{2-\alpha,0}(t)} = r_1 . \tag{17}$$

We argue that this rate is related to the rate observed in FRAP experiments performed on actin networks crosslinked by fluorescently tagged crosslinkers. Unbound linkers diffuse into the bleached region and are responsible for the recovery of fluorescence. Furthermore, detailed balance imposes that the flux of unbinding events from the dark region must equal the flux of binding new fluorescent crosslinkers. As we experimentally find that the diffusion time is much smaller than the unbinding time, we assume that the observed recovery rate equals the unbinding rate of the linkers, $k_{tail}$.

This can be made more explicit by directly considering the dark and fluorescent fractions in the bleached area. Denote by $\pi_i$ the equilibrium fractions in state $i$, and call the dark and fluorescent fractions $p_i^D(t)$ and $p_i^F(t)$ respectively. The fluorescent fraction becomes smaller than $\pi_i$ at $t = 0$, the moment of bleaching. Since the total amount of molecules does not change, we have

$$\pi_i = p_i^D(t) + p_i^F(t) . \tag{18}$$

The assumption of a negligibly small diffusion time makes that $p_0^D(t) = 0$ and $p_0^F(t) = \pi_0$. For the two fluorescent fractions in the bound states the master equation can be given in vector form when we define the vector

$$\mathbf{p}^F(t) = \begin{pmatrix} p_1^F(t) \\ p_2^F(t) \end{pmatrix} . \tag{19}$$

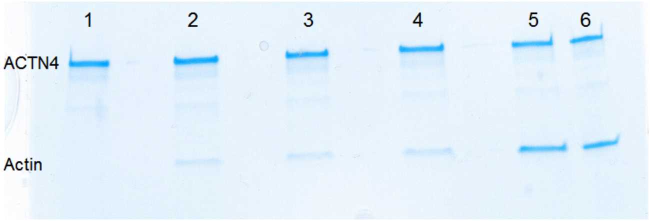


Figure S1. **SDS-PAGE gel showing the biochemical measurement of the binding affinity of alpha-actinin for F-actin.** The supernatant resulting from a high-speed centrifugation of a mixture of actin filaments and crosslinkers was put on a polyacrylamide gel. The top band shows the ACTN4 (100 kDa) and the lower band shows actin (42 kDa). Each channel contains a different actin concentration: 1 = 0 $\mu$M, 2 = 0.2 $\mu$M, 3 = 0.4 $\mu$M, 4 = 0.8 $\mu$M, 5 = 1.6 $\mu$M and 6 = 3.2 $\mu$M.

The master equation is then

$$\begin{aligned} \dot{\mathbf{p}}^F(t) &= \begin{pmatrix} -(k_{10} + k_{12}) & k_{21} \\ k_{12} & -k_{21} \end{pmatrix} \mathbf{p}^F(t) + \begin{pmatrix} k_{01}\pi_0 \\ 0 \end{pmatrix} . && (20) \\ &= -M\mathbf{p}^F(t) + \mathbf{b}. && (21) \end{aligned}$$

In Eq. 21, the negative of the matrix is named $M$ and the constant vector contribution is $\mathbf{b}$. The particular solution of this differential equation is constant,

$$\mathbf{p}_p^F(t) = M^{-1}\mathbf{b}. \tag{22}$$

The homogeneous solutions can be created by finding the eigenvectors and eigenvalues of $M$, and decomposing $\mathbf{p}^F(t)$ into these eigenvectors. The contribution to the solution of each eigenvector decays with a rate equal to its corresponding eigenvalue. Using standard techniques, it can be shown that the eigenvalues of $M$ are exactly $r_1$ and $r_2$. Hence, the combined fraction of fluorescent proteins in the bleached region has a double exponential time dependence with exactly those two rates, until the fluorescence saturates at the steady state value. We can then obtain the asymptotic behavior of the function by taking the long time limit, as done in equation 17. Since $\mathbf{p}^F(t)$ contains the same two exponents as $f_{2-\alpha,0}(t)$, the limit will give the same result, namely that the function is dominated by a single exponential tail for large time, and that the rate equals the slowest rate, in this case $r_1$. We conclude that the fluorescent fraction recovers exponentially with rate $r_1$, when observed at timescales larger than $1/r_2$.

#### Approximation of the recovery rate

For the experimentally relevant regime of small $k_{10}$, the rate in Eq. 12 can be approximated by the mean first passage time [Fig. 6]. First, we look at the general solution Eq. 15 and choose $\alpha$ such that the dark fractions in

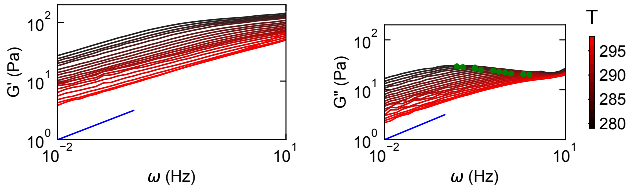


(a) (b)

Figure S2. **Rheology of actin networks crosslinked with alpha-actinin at different temperatures.** Frequency sweeps of the storage (a) and loss (b) shear modulus measured at different temperatures from 280 K (black) to 298 K (red) reveal a decrease of the moduli and a decrease of the characteristic frequency $f_{\mathrm{rheo}}$ (green dots) as a function of temperature (see main text Fig. 2).

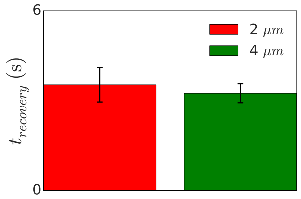


Figure S3. **Effect of FRAP bleach area on the FRAP rate.** The average recovery time for a FRAP radius of 2 $\mu$m is not significantly different from a FRAP radius of 4 $\mu$m, suggesting the recovery time is not significantly affected by the diffusion time of the crosslinkers, but instead is dominated by the crosslinker unbinding time. Data was acquired at 298 K. The error bars represent the standard error on basis of 4 repeats per condition.

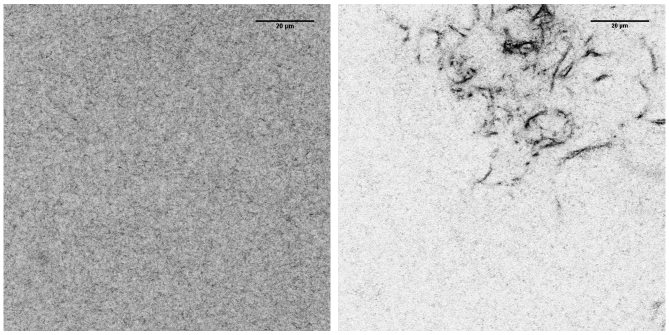

(a) (b)

Figure S4. **Confocal fluorescence images of ACTN4 crosslinked actin networks** a) At a 1:100 ACTN4:actin molar ratio, the concentration used for all experiments in the main text, an isotropically crosslinked actin network is observed with no structure above the diffraction limit. b) For comparison, actin bundles were observed at a 1:25 ACTN4:actin molar ratio. The color coding was inverted for both images to improve the visual contrast between bundles and background. Scale bars are 20 $\mu$m.

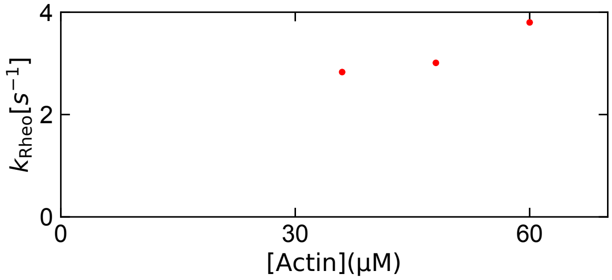


(a)

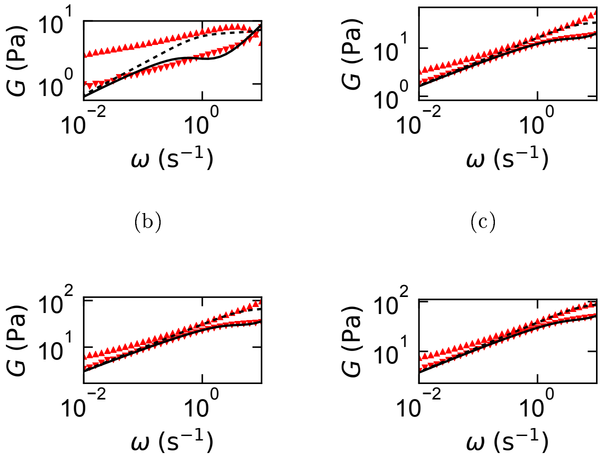


(b) (c) (d) (e)

Figure S5. **Rheology of ACTN4-crosslinked networks at different actin concentrations.** a) In contrast to the FRAP rate, the rate measured by rheology does not decrease with the actin concentration. Instead, a slight increase of the onset frequency of stress relaxation upon an increase of the actin concentration is observed, although that might be due to the imprecision of the fit. Indeed, previous studies found that the timescale measured with rheology is concentration-independent [1–3]. b-e) The storage and loss modulus (upward and downward pointing triangles) as a function of frequency for 24 (b), 36 (c), 48 (d) and 60 $\mu$M actin (e) measured at 298 K. The dashed and solid lines show the fit to eq. 2 in the main text. The timescale measured at an actin concentration of 24 $\mu$M was discarded because of the poor quality of the fit.

state 1 and 2 initially obey detailed balance,

$$\alpha = \frac{k_{21}}{k_{12} + k_{21}}. \tag{23}$$

We call this initial state $\pi$, which gives the first passage time distribution

$$f_{\pi,0}(t) = \frac{k_{10}k_{21}}{r_2 - r_1} \left[ \left(1 - \frac{r_1}{k_{12}+k_{21}}\right) \mathrm{e}^{-r_1 t} + \left(\frac{r_2}{k_{12}+k_{21}} - 1\right) \mathrm{e}^{-r_2 t} \right]. \tag{24}$$

This distribution has mean value

$$\tau_\pi = \frac{k_{12}+k_{21}}{k_{10}k_{21}} + \frac{k_{12}}{k_{21}\left(k_{12}+k_{21}\right)} \approx \frac{k_{12}+k_{21}}{k_{10}k_{21}}. \quad (25)$$

The approximation is valid when $k_{10} \ll k_{12}, k_{21}$. We can approximate Eq. 12 in the same regime, giving

$$r_1 = \frac{1}{2}\left(k_{10}+k_{12}+k_{21}\right)\left(1-\sqrt{1-\frac{4k_{10}k_{21}}{\left(k_{10}+k_{12}+k_{21}\right)^2}}\right), \quad (26)$$

$$\approx \frac{1}{2}\frac{1}{2}\frac{4k_{10}k_{21}}{k_{10}+k_{12}+k_{21}} \quad (27)$$

$$\approx \frac{k_{10}k_{21}}{k_{12}+k_{21}}. \quad (28)$$

The first line uses that $\sqrt{1+\epsilon} \approx 1+\epsilon/2$. Hence, we see that $r_1 \approx 1/\mu_\pi$ when $k_{10}$ is small, and equals $k_{10}$ times the fraction that is in state 1, as given by Eq. 23. The equivalence of approximations through either the asymptotic tail or the mean unbinding time can also be seen in Fig. S6. This means that in our case, the approximation of the rate through a mean first passage time would also have been justified. However, if $k_{10}$ is fast and $k_{12}$ is slow, then the initial fraction in state 1 will drop very rapidly, and the distribution will show a quick drop towards a slow exponential tail with rate $r_1$. In that case, the mean first passage time would get a big influence from the initial drop. Since measurements could very well miss the initial drop because it happens too quickly, the rates cannot be estimated robustly from the mean first passage time. On the other hand, our expression for the asymptotic rate is reliable whenever a single exponential tail is observed in the fluorescence recovery.

##### Equilibrium fractions

The fractions occupying the three states 0, 1, and 2 are set by the transition rates between them and by detailed balance. We call $\pi_i$ the equilibrium, stationary fraction in state $i$. Detailed balance says that

$$k_{01}\pi_0 = k_{10}\pi_1, \quad (29)$$

$$k_{12}\pi_1 = k_{21}\pi_2. \quad (30)$$

Together with the normalization condition $\pi_0+\pi_1+\pi_2 = 1$, this gives the solutions

$$\pi_0 = \frac{k_{10}k_{21}}{k_{10}k_{21}+k_{01}k_{21}+k_{01}k_{12}}, \quad (31)$$

$$\pi_1 = \frac{k_{01}k_{21}}{k_{10}k_{21}+k_{01}k_{21}+k_{01}k_{12}}, \quad (32)$$

$$\pi_2 = \frac{k_{01}k_{12}}{k_{10}k_{21}+k_{01}k_{21}+k_{01}k_{12}}. \quad (33)$$


[1] O. Lieleg, M. M. A. E. Claessens, Y. Luan, and A. R. Bausch. Transient binding and dissipation in cross-linked actin networks. *Physical Review Letters*, 101(10):1–4, 2008.

[2] O. Lieleg, K. M. Schmoller, M. M. A. E. Claessens, and A. R. Bausch. Cytoskeletal polymer networks: Viscoelastic properties are determined by the microscopic interaction potential of cross-links. *Biophysical Journal*, 96(11):4725–4732, 2009.

[3] C. P. Broedersz, M. Depken, N. Y. Yao, M. R. Pollak, D. A. Weitz, and F. C. MacKintosh. Cross-Link-Governed Dynamics of Biopolymer Networks. *Physical Review Letters*, 105(23):238101, nov 2010.

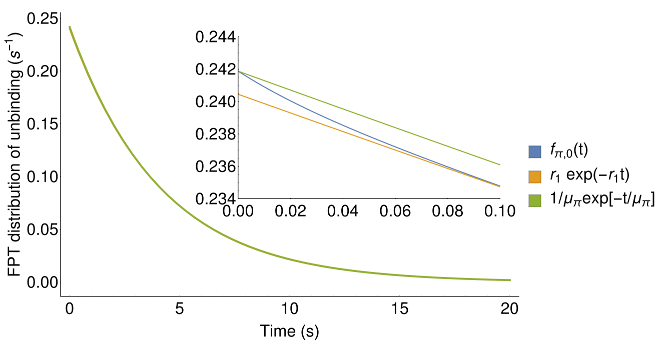


Figure S6. **Crosslinker unbinding kinetics according to the three-state model.** First passage time (FPT) distribution for bound crosslinkers to unbind. Crosslinkers are initially singly or doubly bound with probabilities set by detailed balance, and the FPT distribution gives the probability per unit time that a crosslinker unbinds after that specific time (blue). The exponential distributions with rates equaling the long time tail (orange, Eq. 26) and the approximate mean unbinding time (green, Eq. 25) are also given. For small time, the true distribution deviates slightly from the approximate lines (inset), but all distributions coincide for larger times. We used the experimentally observed parameters of ACTN4 (main text Fig. 3, $k_{10} = 0.43\,\mathrm{s}^{-1}$, $k_{12} = 14\,\mathrm{s}^{-1}$, and $k_{21} = 18\,\mathrm{s}^{-1}$).
